# Homeostasis Back and Forth: An Eco-Evolutionary Perspective of Cancer

**DOI:** 10.1101/092023

**Authors:** David Basanta, Alexander R. A. Anderson

## Abstract

The role of genetic mutations in cancer is indisputable: they are a key source of tumor heterogeneity and drive its evolution to malignancy. But the success of these new mutant cells relies on their ability to disrupt the homeostasis that characterizes healthy tissues. Mutated clones unable to break free from intrinsic and extrinsic homeostatic controls will fail to establish a tumor. Here we will discuss, through the lens of mathematical and computational modeling, why an evolutionary view of cancer needs to be complemented by an ecological perspective in order to understand why cancer cells invade and subsequently transform their environment during progression. Importantly, this ecological perspective needs to account for tissue homeostasis in the organs that tumors invade, since they perturb the normal regulatory dynamics of these tissues, often co-opting them for its own gain. Furthermore, given our current lack of success in treating advanced metastatic cancers through tumor centric therapeutic strategies, we propose that treatments that aim to restore homeostasis could become a promising venue of clinical research. This eco-evolutionary view of cancer requires mechanistic mathematical models in order to both integrate clinical with biological data from different scales but also to detangle the dynamic feedback between the tumor and its environment. Importantly, for these models to be useful, they need to embrace a higher degree of complexity than many mathematical modelers are traditionally comfortable with.

## Introduction

Historically, cancer biology has been an almost entirely empirical field where mathematical theory has been largely neglected. But theoretical frameworks not only allow us to make sense of (often) counter intuitive experimental data but also help define new hypotheses and identify holes in our understanding that need to be addressed (Kuhn 2012). Mathematical models allow us to formally define (and combine) biological mechanisms in such a way that the logical consistency can be checked, the assumptions thoroughly analyzed and scenarios tested at a speed and cost unmatched by experimental models (Möbius & Laan 2015). This is particularly important given the long timescales and multiple interconnected spatial scales involved in the evolution of cancer. Without the aid of mathematical and computational models we have no easy way to understand cancer evolution (Byrne et al. 2006; Servedio et al. 2014).

Mathematical and computational models of cancer have seen rapid growth in the last two decades. The emerging field of Mathematical Oncology, embraces mathematical modelling as a key tool to systematically understand the mechanisms underlying all aspects of cancer progression and treatment and, perhaps even more importantly, to make predictions about future outcomes. In recent years, there have been some excellent reviews on mathematical models of cancer (Anderson & Quaranta, 2008; Byrne 2010; Anon 2014; Korolev et al. 2014; Altrock et al. 2015), our focus here, however, is more explicitly tied to models that consider an ecological view of cancer.

Normal tissues contain a multitude of cell types, molecular signals and microenvironmental features that work in symphony to ensure tissue function, as well as maintain homeostasis. This homeostasis is dynamic and naturally emerges from the interplay between life and the environment (Dyke & Weaver 2013). The majority of cancer models, mathematical or experimental, neglect both the microenvironment in which cancer originates or to which it metastasizes. This is in part due to the dynamic and complex nature of the dialogue between the tumor and its environment. However, since the cancer microenvironment has a strong selective influence on the course of cancer evolution (Mueller & Fusenig 2004; Tlsty 2008; Lu et al. 2012) we cannot continue to ignore it. While genetic mutations can partially explain the diversity that is required for somatic evolution, the physical microenvironment and the interactions with other cells are the source of selection which drives evolution. The importance of the environment in evolution may even justify a new view of evolution based on niche construction which would replace the standard genetic-based view (Turner 2016).

Although normal tissues are robust against many perturbations (Basanta et al. 2008; Gerlee et al. 2011), tumors that will grow to become clinically relevant can disrupt tissue homeostasis beyond the point of recovery. This can lead to a process of clonal evolution (Nowell 1976; Greaves 2015; Gerlinger et al. 2012) where an initial aberrant cell grows, adapts to (and alters) the environment and diversifies. In fact, this diversity known as intratumor heterogeneity (ITH) is a pervasive feature of most cancers and is increasingly correlated with poor prognoses (Gay et al. 2016). There are multiple sources of heterogeneity in cancer (Welch 2016), ITH is not only limited to genetic changes but can include epigenetic (Heng et al. 2009; Angermueller et al. 2016; Easwaran et al. 2014), metabolic (Hensley et al. 2016; Damaghi et al. 2015; Robertson-Tessi et al. 2015), and microenvironmental (Mumenthaler et al. 2015; Natrajan et al. 2016; Lalonde et al. 2014). Importantly this heterogeneity is dynamic (Fisher et al. 2013) which means that where and when mutations occur is equally important (Sottoriva et al. 2015; Swanton 2012). These facts highlight the degree of complexity that makes studying somatic evolution very challenging, especially if we restrict ourselves to experimental techniques alone. Mathematical models can help understand cancer growth (Gatenby & Maini 2003; Michor et al. 2004; Byrne 2010; Anderson & Quaranta 2008; Altrock et al. 2015) and cancer evolutionary dynamics and shed light on the role of ITH in both progression and treatment response (Anderson et al. 2006; Iwasa & Michor 2011).

In this paper we will describe, through the lens of mathematical modelling, how homeostasis disruption can explain cancer initiation, help determine its evolution and suggest novel treatment approaches.

## Models of normal homeostasis and its disruption

The human organism is a complex multiple scale biological system, where multiple organs act as a unified organ system to maintain both health and function. In turn, individual organs are made up of multiple tissues that form specific structures for defined tasks and these tissues contain billions of individual cells that perform diverse functions using many distinct cell types. Much of the communication between individual components of this multiscale system are driven by autocrine and paracrine signals that are either produced by individual cells or as result of interactions between cells. This communication drives a tight dialogue between individual components of the system that regulates and maintains organ and whole organism homeostasis. For example, in the case of bone homeostasis (figures 1 and 4A) the interactions between bone-destroying osteoclasts and bone-producing osteoblasts are mediated by molecules such as TGF-β and RANKL and results in bone remodeling and maintenance. Whereas in the skin (figure 4C–D) keratinocytes (composing 95% of the epidermis) and melanocytes (located on the basement membrane) interact together with factors (EGF, TGFβ and bFGF) secreted by dermal fibroblasts to regulate cell turnover and maintain homeostasis. Homeostasis is also important at the microenvironmental level and key elements like the Extracellular Matrix (ECM) are maintained by a process that, if disrupted, can lead to fibrosis, cancer initiation and metastasis (Cox & Erler 2011).

**Figure 1:**
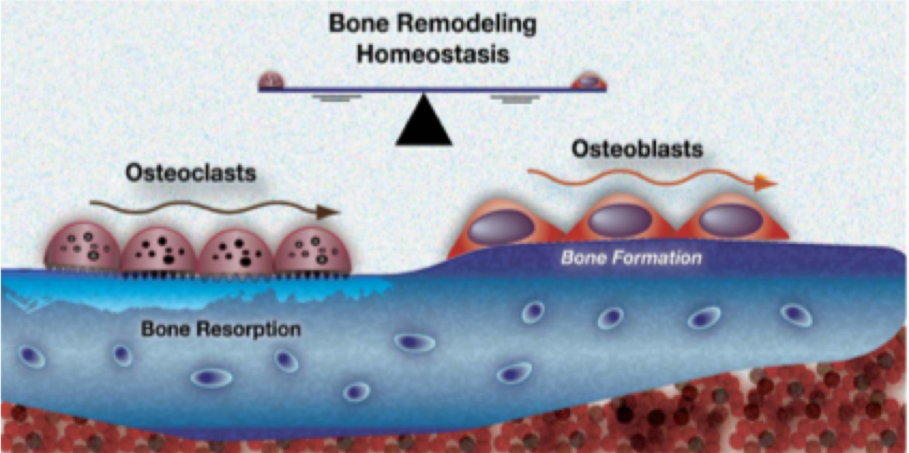
Homeostasis in organs like the bone is dynamic and orchestrated by a variety of cell types and signaling factors. This allows the organ to maintain form and function even in the presence of genetic or environmental insults. Image taken from (Strand et al. 2010).

Homeostasis, at its simplest, is the balance between cell proliferation and apoptosis that preserves the architecture and functionality of the organ. Homeostasis is very robust, since tissues remain fully functional, maintaining shape and size even under significant genetic perturbations and environmental insults.

Organ ecology and homeostasis varies from organ to organ as the mechanisms involved depend on the importance of the organ (Thomas et al. 2016) and the purpose they serve. Mathematical models allow us to explore the mechanisms of homeostasis of relevant tissues, understand how different tumor phenotypes can disrupt this homeostasis, explore how tumor heterogeneity changes through space and time, and understand the impact of new competitive and cooperative interactions on tumor growth and progression. As we discussed previously (Basanta & Anderson 2013) mathematical tools like evolutionary game theory (with its focus on simplicity and cell-cell interactions) and agent-based modelling (with strengths in capturing heterogeneity and the physical microenvironment) are ideal frameworks in which to capture these processes.

We previously used a combination of agent-based modeling and a genetic algorithm to *evolve* homeostatic organisms capable of first growing and then maintaining a target shape and size (Basanta, David et al. 2008). Our results show that evolution of homeostasis also selects for robustness such that the organisms which achieved a higher degree of homeostasis were also better at restoring it after external insult or *genetic* aberrations. In subsequent work (Gerlee et al. 2011) we used a hybrid cellular automaton model to evolve homeostasis and showed that evolution may select for two different strategies to achieve homeostasis: wasteful (with high proliferation and death rates) and conservative (low proliferation and death rates). Both strategies are successful in achieving homeostasis but exhibit different types of susceptibility to mutations which might happen at any stage of the cell’s life cycle or during division. Specifically, wasteful phenotypes were more suited to low mutation rates during replication and higher spontaneous mutation rates (i.e. cosmic ray mutations that can occur at any stage of the cell’s life cycle), whereas the conservative phenotype is better suited to higher mutation rates during replication and low spontaneous mutation rates. Both pieces of research highlight how homeostasis is dynamic and robust to many types of perturbation. More recently Csikasz-Nagy and colleagues have used a combination of game theoretical and network modelling to model homeostasis and how disruption of the interactions between normal cells could help a newly established cancer grow (Csikász-Nagy et al. 2013).

## The Cancer Ecosystem

Tumors are not simply collections of mutated cells that grow in isolation of the environment in which they live. They interact with and modify both the physical microenvironment and a variety of non-tumor cells that make up the organ in which the cancer originated. Given the importance of these interactions it is useful to think of a cancer as an ecosystem, a view that highlights the importance of non-cancer cells and physical microenvironmental features (Merlo et al. 2006; Greaves 2015; Kareva 2011). This view also allows us to better understand cancer initiation, its evolution and the impact of treatment (Pienta et al. 2008; Chen & Pienta 2011; Basanta & Anderson 2013). Perhaps the most important, but usually overlooked, feature of organ ecosystems is that before cancer developed the organ was a dynamic and functional homeostatic system. An ecology view of cancer envisions tumor cells as a new species invading a healthy ecosystem disrupting its homeostasis (Lloyd et al. 2016). Not all invading species are successful but those that are, can take advantage of their local environment as well as find weaknesses in the mechanisms that control homeostasis. Thus homeostatic mechanisms not only constitute a barrier that needs to be overcome by an evolving tumor, but also means that understanding them is the first step in understanding the evolutionary trajectory of a tumor (Frank & Rosner 2012). This is especially important considering that early mutations might have a disproportionate impact on how tumors evolve (Sottoriva et al. 2015).

Like many cancers, prostate cancer becomes lethal when it metastasizes and, in 90% of the patients there is evidence of metastases to the bone. The elements that make up the bone ecosystem that prostate cancer might subsequently disrupt are well known (Mundy 2002). Figure 2 shows some of the elements that characterize the normal bone ecosystem (left) and how a successful (and heterogeneous) metastasis can disrupt that homeostasis leading to a vicious cycle promoting metastatic growth (center). Mathematical models (Araujo et al. 2014) have allowed us to explore what phenotypes usually lead to successful metastases.

**Figure 2:**
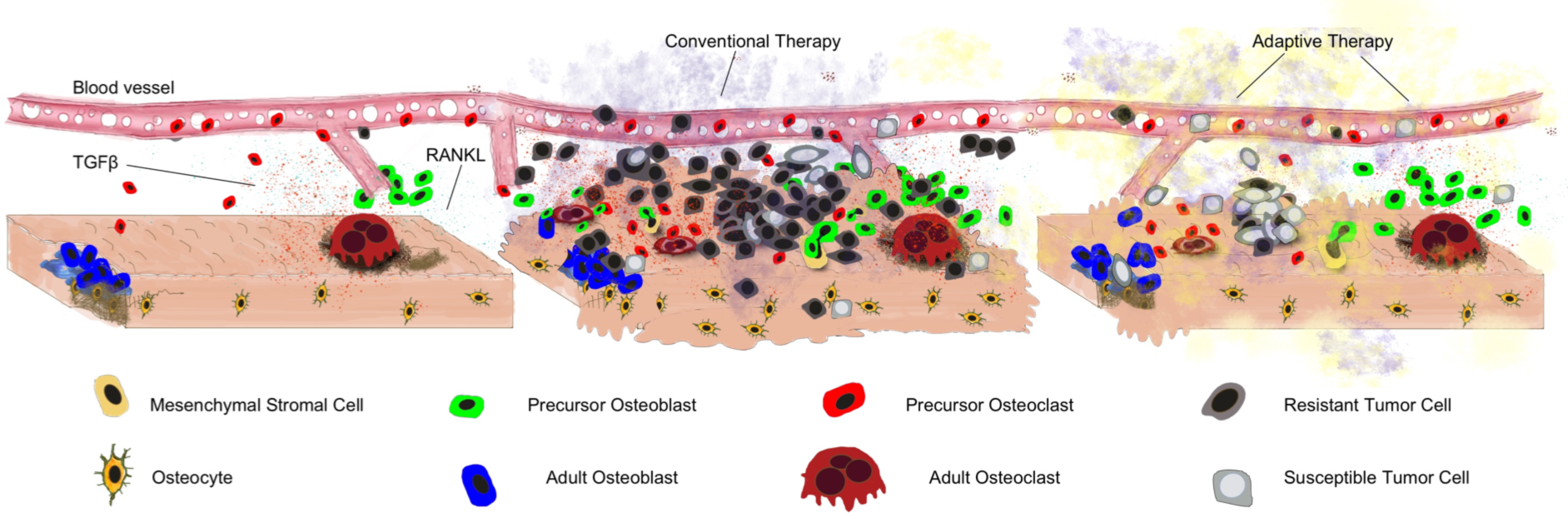
Tissue Homeostasis, Disruption and Restoration. An example of dynamic tissue homeostasis in the bone including bone resorpting osteoclasts and osteoblasts (left). With the introduction of tumor cells (centre), this homeostasis is disrupted leading to an altered microenvironment and increasingly malignant tumor. Conventional application of treatments leads to an increasingly resistant tumor. (right) Eco-evolutionary enlightened treatments aim to restore some degree of homeostasis while allowing the tumor to remain treatable.

## Evolution and Selection

It is well known that selection acts on the phenotype, not the genotype. Hanahan and Weinberg summarized decades of research in cancer genetics to identify six, later extended to ten, phenotypic hallmarks that characterize the emergence of aggressive metastatic cancers (Hanahan & Weinberg 2011). Axelrod and colleagues suggested that, given the right ITH, a group of cancer cells could collectively gain all the hallmarks to be an aggressive metastatic cancer (Axelrod, Axelrod & Pienta 2006b). While it is assumed that competition is the main type of interaction between cancer cells in a tumor, cooperative behavior could help the tumor achieve all the hallmarks much faster than in the case in which every cell has to gain all the phenotypic traits described by Hanahan and Weinberg. Together with spatial heterogeneity (Komarova 2014; Kaznatcheev et al. 2015), the ability of the tumor to evolve is limited by phenotypic heterogeneity (Korolev et al. 2014; McGranahan & Swanton 2015a). It is clear then that, although we are now beginning to understand some of the mechanisms that underpin heterogeneity (Sutherland & Visvader 2015) and how that heterogeneity correlates with prognosis (Gay et al. 2016) and treatment response (Junttila & de Sauvage 2013), its non-trivial to properly understand how evolution impacts cancer progression (Miller et al. 1989) which further emphasizes the need for mathematical modelling.

Mathematical tools like evolutionary game theory (Smith 1982; Basanta & Deutsch 2008; Hummert et al. 2014) can be used to investigate how the interactions between tumor cells, with different phenotypic strategies, can impact selection (Basanta 2015). Tomlinson investigated angiogenesis as a public good (Tomlinson & Bodmer 1997), work that was followed by Bach and colleagues (Bach et al. 2006) examining how an effort that requires cooperation (such as angiogenesis) can emerge in a tumor population. The results in both models show that this type of cooperation would allow for one of the cancer hallmarks to emerge even when many tumor cells are non-angiogenic. Further work by Archetti and colleagues, using evolutionary game theory and *in vitro* modeling, explains how cooperation and ITH can be maintained with other factors like IGF (a growth factor) that can effectively be considered as public goods (Archetti et al. 2015).

Evolutionary game theory can provide further insights on the interplay between heterogeneity and evolution. For instance, we explored the emergence of invasive phenotypes in cancer using a simple game between two tumor cell phenotypes, purely proliferative and motile cells (Basanta et al. 2008) (figure 3). In this game the only factor driving the emergence of invasive phenotypes was the cost of motility (which depends on the type of cancer such that the cost is lower in cancers like leukemia and higher in solid tumors). The introduction of a new phenotypic strategy, that of tumor cells with a glycolytic metabolism (resulting in acidification of the microenvironment leads to selection for invasive phenotypes when we consider the scenarios that, paradoxically, should favor glycolytic cells (e.g. when the cost of environmental acidification is higher for proliferative cells). Even if glycolytic cells were not abundant in the game, the results show that they played a key role in the evolutionary dynamics of the cancer in ways that would be hard to elucidate with experimental data alone.

**Figure 3:**
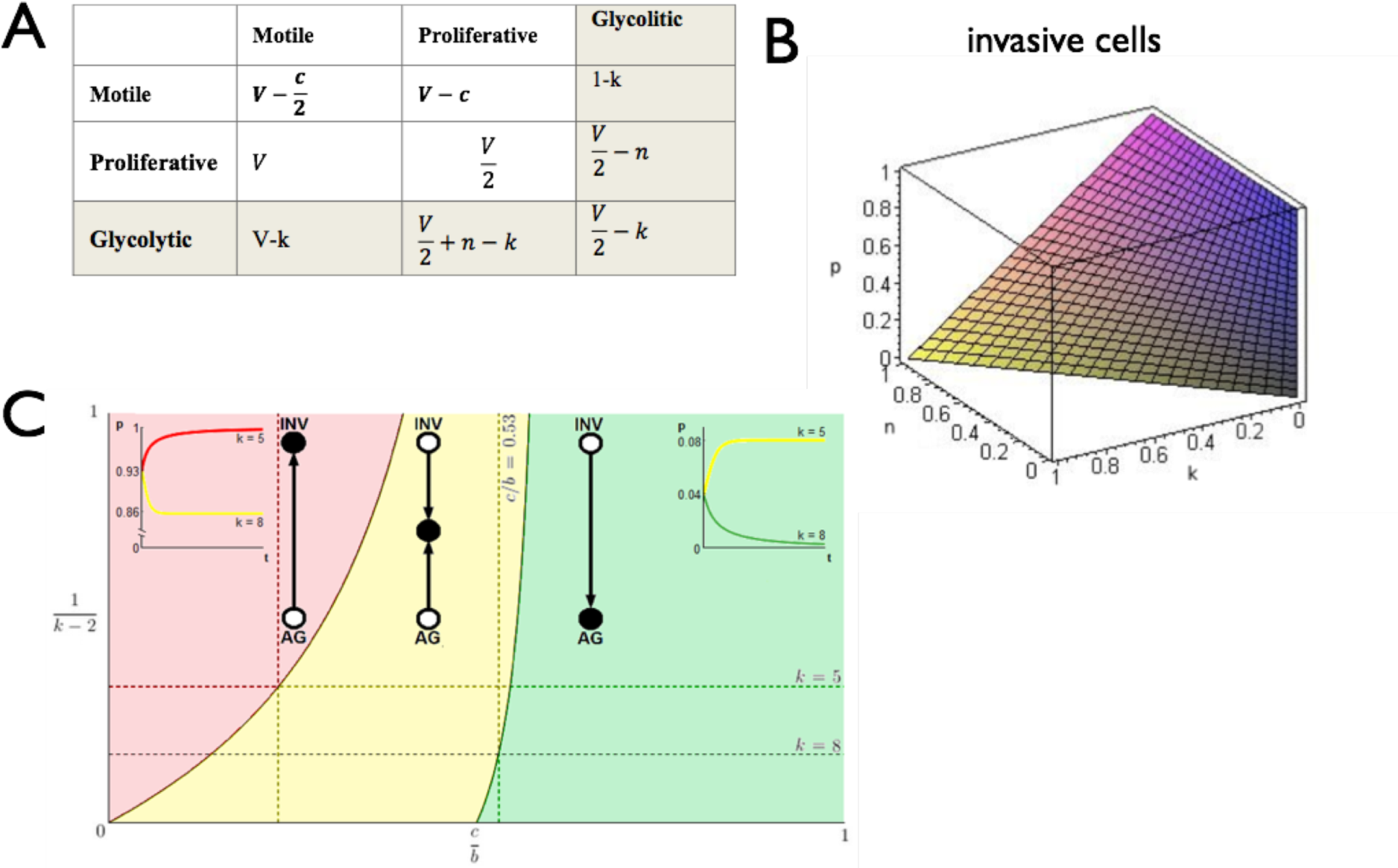
Game Theory is a tool to understand the impact of interactions in selection. (A) Payoff table describing the interactions between proliferative, invasive and glycolytic cells. (B) Proportion of invasive cells goes up as we increase the heterogeneity of the original game to consider glycolytic cells. (C) Space also plays a role in selection, using an Othsuki-Nowak transform we investigated the impact of hard edges (like bone) upon evolutionary dynamics. Here, we plot level of viscosity 1/(k-2) versus relative cost of motility c/b and highlight three regions with qualitatively different dynamics. In the red, the population evolves towards all invasive (INV); in the yellow—towards a polyclonal tumour of INV and proliferative cells (AG); and in the green the tumour remains all AG.

Evolution is driven by stochastic processes and is notoriously difficult to predict (Lipinski et al. 2016), this is especially true for cancer, even with the help of a mathematical model. Although, this does not mean cancer evolution is entirely stochastic. There are constraints that could help define potential evolutionary trajectories (Venkatesan & Swanton 2016). Some of these constraints are related to the specific location of the tumor cells with regards to the rest of the cancer (Lloyd et al. 2016) and homeostasis itself may dictate which intrinsic and extrinsic barriers need to be broken first. Therefore, by understanding the microenvironmental context in which tumors emerge and the homeostatic mechanisms that need to be disrupted (Gerlee et al. 2011; Araujo et al. 2014) we can better understand the evolutionary trajectories of successful tumor cells. As Hanahan and Weinberg’s work shows, while the specific genetic mutations might not be clear, what is clear are the traits that need to be acquired by a successful cancer initiating cell or metastasis. In the case of prostate cancer to bone metastases for instance, being able to disrupt the dialogue between bone-resorbing osteoclasts and bone-producing osteoblasts. This is usually accomplished through the upregulation of the signaling molecule TGF-β. Therefore, understanding the mechanisms that regulate tissue homeostasis will allow us to better predict the first steps in the evolutionary process, which in certain cancers could be the only steps that matter (Sottoriva et al. 2015).

In some tumors, it has been observed that mutations which drive evolution are present from the early stages (Sottoriva et al. 2015) restricting the number of evolutionary paths in subsequent progression i.e. first versus fittest (Robertson-Tessi & Anderson 2015). Even with the limitations of commonly used sequencing tools, such as Sanger sequencing (Schmitt et al. 2016), they are currently the best way to quantify ITH (McGranahan & Swanton 2015a). Bioinformatic tools allow for the extensive interrogation of molecular data to uncover driver mutations (Gonzalez-Perez, Perez-Llamas et al. 2013, Zhang, Liu et al. 2014, Forbes, Beare et al. 2015), molecular signatures that characterize a variety of cancer types and evolutionary paths (Nielsen 2005, Sotiriou & Piccart 2007, Liberzon et al. 2011, Alexandrov et al. 2013; Alexandrov & Stratton 2014) and even potential immunogenicity (Snyder and Chan 2015). But this analysis is fundamentally correlative, tumor centric and provides only a cursory understanding of the underlying biological mechanisms at play without exhaustive substantiation. However, we do not mean to imply they are without utility, in fact there is an opportunity to integrate what we learn from the molecular scale with data from the tissue (histology) and organ (imaging) scales through mechanistic mathematical models that embrace the microenvironment. This type of integrated analysis and modelling is still in its infancy but it has great potential to bridge the divide between the data rich molecular and data poor clinical aspects of cancer (Sottoriva et al. 2015).

## Eco-Evolutionary Models of Cancer

Given the importance of homeostasis in regulating tissue form and function it is surprising that models (experimental and mathematical) typically assume that tumors are present at the start, have already disrupted tissue homeostasis and ignore surrounding tissue interactions. The necessity to disrupt homeostasis represents a key adaptation for tumor cells and, in complex non-linear systems, this transition can be sudden and difficult to observe experimentally (Trefois et al. 2015). Importantly, understanding homeostatic mechanisms can help predict robustness (Scheffer et al. 2012), identify tipping points (Scheffer et al. 2009) and anticipate the most likely evolutionary dynamics of the runaway cancer. In this section we will discuss some models that consider normal tissue homeostasis as a prerequisite for tumor initiation. This invariably means the models are more complex and often multiscale, since they need to take into account both tumor centric and environment centric mechanisms as well as the dialogue between them.

Hybrid models provide a natural way to integrate both discrete and continuous variables that are used to represent individual cells and microenvironmental variables, respectively (Rejniak & Anderson 2011). Each discrete cell can also be equipped with submodels that drive cell behavior in response to microenvironmental cues. Moreover, the individual cells can interact with one another to form and act as an integrated tissue. In our previous work we employed such a hybrid cellular automaton (HCA) approach to characterize the glandular architecture of prostate tissue and its homeostasis through a layered epithelial homeostasis via TGF-Beta signaling regulated by surrounding stroma (Basanta et al. 2009). Using this model we examined (see figure 4B) prostate cancer initiation and were able to capture prostatic intraepithelial neoplasia (PIN) as well as invasion and make some interesting hypotheses regarding the differential effects of stroma and TGF-Beta. For example in PIN, TGF-Beta recruits stromal cells, which structurally inhibit glandular breakdown through matrix production, whereas TGF-Beta is more likely to promote growth once tumor cells emerge from a contained PIN-like state and facilitate stromal activation. In subsequent work we used a similar computational approach to explore bone homeostasis emerging from the interactions between osteoclasts, osteoblasts and other bone-resident cells (Araujo et al. 2014). Using this model, we investigated how prostate cancer cells can disrupt bone remodeling and how this disruption leads to a vicious cycle that enables the tumor to grow.

**Figure 4:**
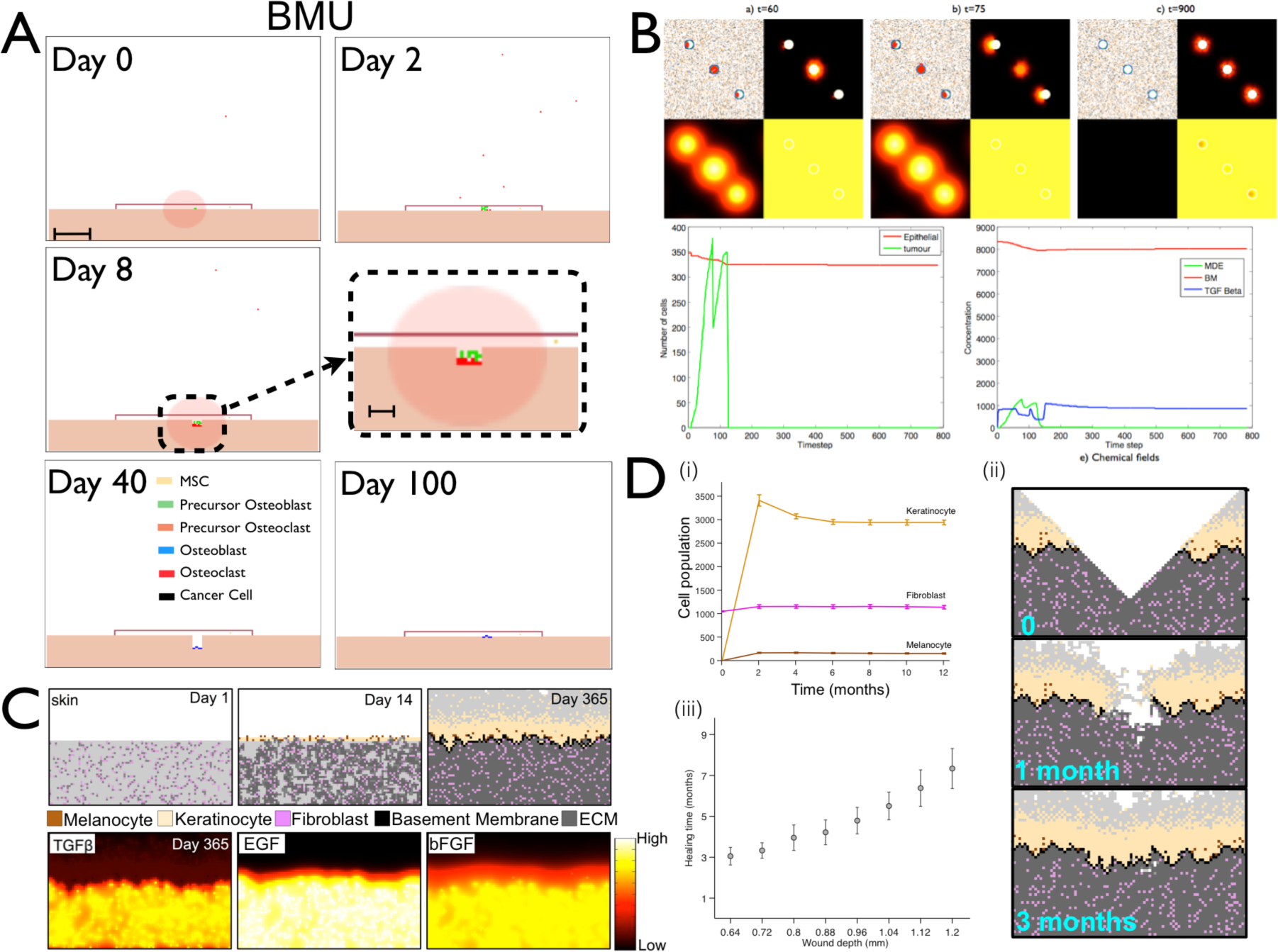
(A) Agent-based models such as the HCA can describe complex homeostasis like the one in the bone, orchestrated by several cell types and involving a number of signaling molecules (Araujo et al. 2014). (B) An HCA model of prostate cancer with homeostasis of the epithelial cells. While not always the case, in some simulations homeostasis can be restored even after cancer initiation. (C) An HCA model of melanoma initiation, showing a homeostatic layer of skin growing over the course of a year via interactions between 3 cell populations (melanocytes, keratinocytes and fibroblasts) modulated by growth factors (TGFb, EGF and bFGF). (D) (i) Cell population dynamics highlight homeostatic fractions; (ii) An emergent property of the homeostatic skin structure is its ability to heal in response to wounding events of differing depths (iii).

In order to understand melanoma initiation we developed an in silico model of normal skin, that incorporates keratinocytes, melanocytes, fibroblasts and other key microenvironmental components of the skin (Kim et al. 2015). The model recapitulates normal skin structure that is robust enough to withstand physical (Figure 4C,D) as well as biochemical perturbations. An important prediction this model generated was that senescent fibroblasts can create a favorable environment for melanoma initiation by facilitating mutant melanocyte growth through growth factor production and matrix degradation. This suggests that senescent fibroblasts should be considered as a potential therapeutic target in the early stages of melanoma progression.

In a recent paper we developed an HCA model to investigate the evolution of acid-mediated invasion (Robertson-Tessi et al. 2015). The model incorporates normal cells, blood vessels, aerobic, glycolytic, acid resistant tumor cells, as well as environmental variables such as oxygen, glucose and acidosis. Tumor cells in the model have two continuously variable, heritable traits: glucose consumption and resistance to extracellular pH (Fig. 5a). Cells interact through the HCA decision algorithm (Fig.5b). Importantly, the normal cells and blood vessels, along with cell oxygen, glucose production/consumption form a homeostatic tissue. An emergent property of this model is that normal tissue can recover from both chemical and physical damage. The model examines the evolution of tumour phenotypes over time into this normal tissue space, finding that poor vasculature leads to acid-resistance in the center of the tumor, followed by selection for aggressively glycolytic cells (Fig. 5c–f). Eventually, these metabolically atypical clones escape the niche that selected for them and interface with the normal tissue. The excess acidosis created by these cells destroys normal tissue and leads to rapid invasion (Fig. 5g,h). Varying the vessel density in time and space leads to different patterns of necrosis and invasive phenotypic distributions, a possible explanation for different histological patters seen in different grades of cancers. This model also highlighted the risks of cytotoxic and antiangiogenic treatments in the context of tumor heterogeneity resulting from a selection for more aggressive behaviors.

**Figure. 5:**
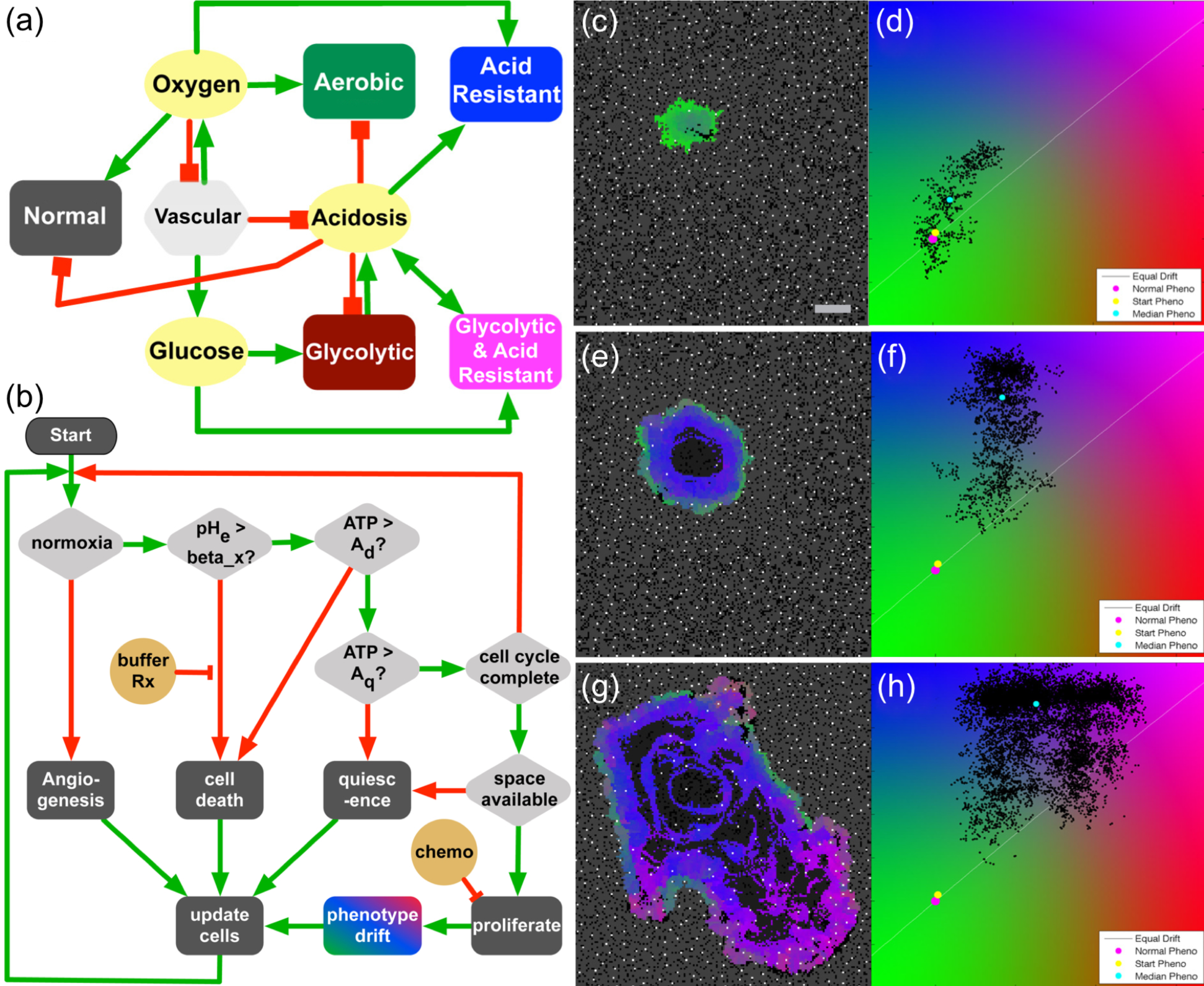
Multiscale metabolic cancer invasion model. (a) HCA model Interaction network highlights positive (green) and negative (red) cellular and microenvironmental interactions. (b) Cell life cycle flowchart that every cell experiences, dictating cell proliferation, death and mutation depending on the levels of ATP, Oxygen and pH. (c,e,g) Heterogeneous tumor growth and evolution at 2, 8 and 20 weeks. (d,f,h) Corresponding phenotypic (x-axis=glycolytic activity-axis=acid resistance) distribution at the same simulation time points.

## Bridging the Divide

Clinical trials are the standard way in which basic science discoveries are translated into clinical practice. A major issue with these trials in cancer is their high failure rate, even though the trials were driven by data from successful equivalent preclinical studies. This is in part due to the intrinsic homogeneity of preclinical model systems and the contrasting heterogeneity of actual patient responses. It is becoming increasingly clear that not all pre-clinical outcomes scale to human populations and that mathematical models need to be developed with this in mind to allow researchers to explore how intra patient heterogeneity in human cohorts would impact novel therapies developed and tested using *in vitro* and/or *in vivo* models (Kim et al. 2016).

We and others have used clinically and experimentally parameterized mathematical models to predict and to understand the impact of treatments in cancer. Mechanistic first-principles mathematical models help us not only to predict **when** treatments fail to work but also help us understand **how** they fail. However, this understanding has largely been focused on a tumor centric perspective and whilst this is certainly important, we only have a very limited understanding of how treatments impact normal cells and the microenvironment of the tumor (Medler et al. 2015). This further highlights the importance of mathematical models and how they could be used to inform clinical trials (Rockne et al. 2010; Mumenthaler et al. 2011; Scott 2012; Leder et al. 2014; Gallaher et al. 2014; Baldock et al. 2014; Walker & Enderling 2015; Kim et al. 2015; Poleszczuk et al. 2016).

These mathematical models can help us better understand how interventions impacting key genetic or epigenetic drivers, may or may not shape the evolutionary dynamics of the disease in the right direction. But for these models to be useful, and to understand homeostatic mechanisms, they need to be able to account for the interactions between cells and between those cells and their environment. That is why network models, evolutionary game theory (Basanta 2015; Brown 2016) and especially hybrid agent-based models that explicitly account for environment and heterogeneity are so important (Anderson 2005; Anderson et al. 2006; Gerlee & Anderson 2008;).

Hybrid agent-based models can also be used to investigate the impact of targeting specific aspects of the tumor system. Often, what looks like a key driver of an invasive process that can be therapeutically targeted but might result in a tumor that is more invasive or difficult to treat. For instance, in bone metastatic prostate cancer stromal cells like osteoclasts can be targeted by treatments like bisphosphonates but experimental (Logothetis & Lin 2005) and mathematical research (Basanta et al. 2011; Araujo et al. 2014) show that this approach is not sufficient to prevent successful metastatic growth. In other cases, treatments that have shown poor results in the clinic can be rejuvenated through different dosing or combination with others. Well characterized molecules such as TGF-β for instance might play dual roles (Bierie & Moses 2006) but only through careful mathematical modeling are we be able to understand whether they present clear clinical targets (Basanta et al. 2009; Cook et al. 2016). In the multiscale metabolic model we discussed in figure 5 above, targeting the vasculature through an anti-angiogenic therapy leads to a more aggressive and invasive cancer (Robertson-Tessi et al. 2015). This is because the therapy actually induces hypoxia and accelerates the acquisition of the acid-resistant/glycolytic phenotype, which may be a key reason for the failure of many anti-angiogenic therapies (Rapisarda & Melillo 2012; Quintieri et al. 2014).

## Adaptive Therapy and Homeostasis Restoration

In the context of cancer as an evolutionary process it is sometimes easy to forget that treatments constitute selection. The prevalence of intra-tumor heterogeneity means that some clones will be negatively impacted by treatment but also that some will be left unaffected by it. Although treatment can reduce tumor burden and also, even if transiently, reduce heterogeneity (Gerlinger & Swanton 2010), the emergence of resistance is common. Relapse is usually driven by clones that are not the linear descendent of the most common clone at diagnosis (Wang et al. 2016). We need to understand how treatment-derived selection facilitates resistance (Hata et al. 2016) and changes the tumor, in ways that make further treatment more difficult if not impossible e.g. multidrug resistance. Critical to this understanding are the costs that treatments represent to the different phenotypic strategies present in an evolving tumor. Since this will allow us to synergize treatments such that evolving resistance to one drug makes cells more susceptible to another (Orlando et al. 2012; Basanta et al. 2011; Basanta et al. 2012). Traditionally, cancer cells have been seen as competing with each other but in reality a variety of interactions could be taking place within a cancer (Axelrod, Axelrod & Pienta 2006b; Strand et al. 2010; Thomas et al. 2012; Basanta & Anderson 2013; Brown 2016). Cooperative behaviors allow tumor cells to acquire cancer hallmarks faster but also suggest that disrupting this cooperation could seriously impact the tumor’s overall aggressiveness (R. Axelrod, D. E. Axelrod & Pienta 2006a; McGranahan & Swanton 2015b; Archetti 2013). Targeted treatments have been less successful than initially anticipated, in part due to their continuous delivery which inevitably leads to selection for resistance (Gillies et al. 2012). However, if used in combination with other treatments, through the lens of evolution they could help to steer the evolutionary dynamics of the disease to facilitate long term control.

Until recently, most cancer treatments aimed to kill as many tumor cells as possible while minimizing damage to healthy ones. This approach does not take into consideration heterogeneity (Alfonso et al. 2014) or how they impact selection in somatic evolution (Greaves & Maley 2012; Miller et al. 1989; Gerlinger & Swanton 2010). Evolutionary enlightened therapies such as adaptive therapy (figure 6) constitute one important step towards treatments that not only understand this impact but also exploit it. The theoretical framework underlying adaptive therapies (Gatenby et al. 2009) has been tested experimentally (Enriquez-Navas et al. 2016) and is now being used in a clinical trial at the Moffitt Cancer Center (NCT02415621). Gatenby and colleagues recognized that the standard maximum tolerated dose approach used in many treatments, kills all the sensitive cells to allow for unopposed proliferation of any remaining resistant cells, this phenomenon is called competitive release (Adkins & Shabbir 2014). Their adaptive strategy instead is to design treatments that aim for control by leaving behind a residual population of drug sensitive cells, rather than only resistant ones, which will regrow when treatment is paused and allow for subsequent future treatment applications. This ultimately means each patient’s treatment will be adapted based on their response rather than any single fixed regime. This effort is likely to be only the beginning of other mathematically-led efforts with not only biological but also clinical impact. Subsequent work should look not only at the competitive interactions between resistant and susceptible tumor cells but also at the myriad of interactions between a heterogeneous tumor with stromal and immune cells (Barron & Rowley 2012) as well as the microenvironment (Meades et al. 2009; Hirata et al. 2015; Fedorenko & Smalley 2015).

Furthermore, an understanding of how treatments differentially impact tumor cells (Gao et al. 2015; Andor et al. 2015) and the microenvironment would allow mathematical models to play the long game: to use treatments not as ends in themselves but as means to steer somatic evolution in a direction where the tumor either finds itself, in an evolutionary dead-end (Maley et al. 2004), its growth can be controlled for long periods of time (Gatenby et al. 2009) or its size reduced permanently (Foo & Michor 2009; Gallaher et al. 2014). Halting evolution once it has started is incredibly difficult (Greaves 2015; Sottoriva et al. 2015; Robertson-Tessi & Anderson 2015) but steering it might be an option for the next generation of oncologists with sufficient treatment options and time to deploy them. Critically all this information needs to be integrated into mathematical/computational frameworks in a way that is patient-specific - any steering or anti-evolutionary treatment strategy must be tailored to a specific cancer in a given patient, which inevitably means the end of fixed treatment regimens.

Together these facts suggest that in the future we will implement clinical approaches that shape the evolutionary dynamics of tumors (rather than ignoring them) making them easier to treat, with fewer doses of drug and therefore reducing the impact on quality of life. It is possible that this might lead to a sequence of treatment combinations that completely eradicate the tumor. However, the lack of adequate targeted treatments or the presence of resistant clones that cannot be impacted either directly or indirectly (including other cell types or factors supporting their growth or resistance) means that we should focus on control rather than eradication, especially for disseminated disease. Adaptive Therapies as championed by Gatenby (Gatenby et al. 2009) represent a substantial step in this direction. Although not presented this way, adaptive therapies constitute an effort to accept that tumor cells cannot be eradicated and thus some sort of compromise needs to be enforced where homeostasis is the goal. As illustrated in figure 2 (left and centre), tumors become a disease because they disrupt homeostasis. Novel treatment strategies could be designed and guided by mathematical and computational models (Gallaher et al. 2014) to steer evolution towards a return to homeostasis, even if different from the normal tissue before carcinogenesis (figure 2, right). This treatment-enforced homeostasis could give new hope to patients with the more heterogeneous and disseminated cancers, a hope that existing treatment strategies do not offer.

## A Delicate Balance Between Complexity and Understanding

The biological community has long acknowledged the complexity of cancer and has developed a dizzying array of tools to dissect, quantify and test almost every aspect of it through all stages of progression and treatment. This reductionist paradigm has given us a detailed understanding of many of the component parts that define cancer as exemplified by the seminal Hallmarks papers (Hanahan & Weinberg 2000, 2011). However, we are now left with the fundamental problem of how to piece these individual components back together to define the cancer system. The enormous data sets already collated and currently being generated, largely at the molecular scale, need to not only be mined for correlations but also to be integrated with data across biological scales to understand causation. Bridging the divide between genotype and phenotype, between cells and tissues, organs and organ systems, individuals and populations remains an incredibly challenging and understudied question. However, this question is critical to the understanding and future success of cancer research and why integrated mathematical/experimental/clinical approaches are so important.

Taken together, the reductionist component parts and the multiscale nature of cancer, its clear we have an overwhelming menu of potential players in building eco-evolutionary models of cancer – this begs the question what should we include and what should we leave out? One might be tempted to try to include everything and build the ultimate ’kitchen sink’ model of cancer. But building a model that is almost as complex as the real system would only provide a poor cartoon version of reality that may mimic it but provide little gain in our understanding of how it works. Aiming for truly minimal models is not an option if we aim, as this piece advocates, to understand key aspects of cancer progression such as homeostasis disruption. A sufficient degree of realism and complexity is required to avoid biasing the evolutionary route that a tumor might take to that hard-coded into the model. Therefore, we need to be cautious and build models that walk the tightrope between complexity and understanding, to include sufficient details as to be useful and critically understandable.

Ultimately, this highlights the fact that building mathematical models of cancer is somewhat of an art that requires skill, thought, intuition, a delicate balance between complexity and simplicity and, crucially, a dialogue with biologists. Placing these important variables in the right modelling framework can produce novel insights into the fundamentals of the cancer process and naturally lead to experimentally testable hypotheses. Therefore, it is clear that the key lies really in the biological question that needs to be answered. This should drive the model derivation and define the scales at which the model operates and bridges. In the context of this review, eco-evolutionary models will always need to be more complex since they integrate both tumor and microenvironment aspects of cancer and are driven by the central premise of developing in normal homeostatic systems. However this complexity, by constraining model dynamics, helps the model better forecast evolution and avoid dynamics that, while interesting, will have little relevance to those observed in biology or in the clinic. We believe that modular approaches to integrate multiple mathematical models into a more complex eco-evolutionary system will become one of the more important future directions in the mathematical oncology community.

## Acknowledgements

We would like to thank Dr. Arturo Araujo for his help in producing the figures in this manuscript. We would also like to thank Dr. Robert Gatenby for helpful discussions regarding adaptive therapies. Finally, DB acknowledges a Moffitt Team Science award and grant NCI U01CA202958-01 and DB and ARA acknowledge NCI U01CA151924 and ARA acknowledges NCI U54CA193489.

